# Microengineering 3D Collagen Matrices with Tumor-Mimetic Gradients in Fiber Alignment

**DOI:** 10.1101/2023.07.09.548253

**Authors:** Indranil M. Joshi, Mehran Mansouri, Adeel Ahmed, Richard A. Simon, Poorya Esmaili Bambizi, Danielle E. Desa, Tresa M. Elias, Edward B. Brown, Vinay V. Abhyankar

## Abstract

In the tumor microenvironment (TME), collagen fibers facilitate tumor cell migration through the extracellular matrix. Previous studies have focused on studying the responses of cells on uniformly aligned or randomly aligned collagen fibers. However, the in vivo environment also features spatial gradients in alignment, which arise from the local reorganization of the matrix architecture due to cell-induced traction forces. Although there has been extensive research on how cells respond to graded biophysical cues, such as stiffness, porosity, and ligand density, the cellular responses to physiological fiber alignment gradients have been largely unexplored. This is due, in part, to a lack of robust experimental techniques to create controlled alignment gradients in natural materials. In this study, we image tumor biopsy samples and characterize the alignment gradients present in the TME. To replicate physiological gradients, we introduce a first-of-its-kind biofabrication technique that utilizes a microfluidic channel with constricting and expanding geometry to engineer 3D collagen hydrogels with tunable fiber alignment gradients that range from sub-millimeter to millimeter length scales. Our modular approach allows easy access to the microengineered gradient gels, and we demonstrate that HUVECs migrate in response to the fiber architecture. We provide preliminary evidence suggesting that MDA-MB-231 cell aggregates, patterned onto a specific location on the alignment gradient, exhibit preferential migration towards increasing alignment. This finding suggests that alignment gradients could serve as an additional taxis cue in the ECM. Importantly, our study represents the first successful engineering of continuous gradients of fiber alignment in soft, natural materials. We anticipate that our user-friendly platform, which needs no specialized equipment, will offer new experimental capabilities to study the impact of fiber-based contact guidance on directed cell migration.

## 1. Introduction

Type I collagen fibers are crucial in coordinating cellular activities in native tissues, with aligned fiber domains extending over sub-millimeter to millimeter length scales[1–3]. At the millimeter scale, fiber alignment maintains the organization of cardiac, skeletal, and smooth muscle cells and enables directional force generation along the axis of alignment [4–6]. At the sub-millimeter length scale, aligned fibers guide epithelial, endothelial, and immune cells during tumor invasion[7,8], angiogenesis[9,10], and immune trafficking processes[11], respectively. Here, the aligned fibers provide biophysical contact guidance cues that help cells navigate toward or away from vascular and lymph vessels during the matrix invasion phase of the metastatic cascade. Indeed, in biopsy samples of breast[7], ovarian[12], and pancreatic cancer[8], collagen fibers oriented perpendicular to the tumor interface are a clinical predictor of metastases and poor patient outcomes. Over the past decade, the biological significance of fiber alignment has led to the development of in vitro platforms to study the impact of ECM architecture on cell behavior in healthy and diseased tissue[13,14].

The overarching goal of 3D in vitro model systems is to improve the physiological relevance of experiments by replicating tissue-specific environmental cues and matrix architectures. To this end, collagen is a popular biofabrication material because it is the most abundant structural protein in the body and can create hydrogel scaffolds with defined fiber alignment landscapes in the laboratory. As reviewed by several groups, collagen fibers can be aligned using bioprinting approaches[15,16], electrospinning [17,18], or by applying external magnetic[19], electric, strain[20,21], and flow fields[22,23]. Alternatively, collagen fibers can be reorganized and aligned using embedded, contractile cells followed by decellularization to produce a hydrogel with uniform fiber alignment[14,24,25]. Although these methods effectively create collagen-based hydrogels with homogeneous, millimeter-scale fiber alignment, they cannot replicate the more complex sub-millimeter scale alignment found in vivo.

The ECM can exhibit sub-millimeter spatial heterogeneity because the matrix architecture is actively reorganized in response to cell-induced traction forces[26]. These cell-substrate interactions introduce local gradients in matrix properties that are sensed and interpreted by resident cells. For example, the motility of epithelial, endothelial, and immune cells in response to graded biophysical “taxis” cues, including stiffness (durotaxis)[27,28], topography (topotaxis)[29,30], tethered biomolecules (haptotaxis)[31], have been extensively studied. Importantly, there is evidence that cells respond to spatially graded cues differently than homogeneous signals[32–34]. This phenomenon is particularly relevant to the tumor microenvironment (TME), where cell-matrix interactions can introduce sub-millimeter fiber alignment landscapes that decay with distance away from the interface (i.e., fiber alignment gradients)[35]. However, studies that explore the role of contact guidance in natural materials are primarily limited to collagen or fibrin hydrogels with uniformly aligned fiber architectures[14,36]. Thus, even though the importance of graded biophysical cues is well-recognized, fiber alignment gradients have not been widely investigated as a potential guidance cue, likely because an experimentally tractable method to engineer complex alignment gradients in natural materials has not been available.

To help develop increasingly biologically relevant in vitro models, we present a geometry-based approach using a constricting and expanding channel design to engineer 3D collagen gels with continuous, graded fiber alignment spanning sub-millimeter to millimeter length scales that can be tuned simply by adjusting the input flow rate. We also describe a flexible experimental approach that enables cells and cell aggregates to be patterned at defined locations on the alignment gradient. As a proof of concept, we show that human umbilical vein endothelial cells (HUVECs) migrate directionally on gradient gels compared to random gels and present preliminary evidence that local alignment gradients can act as guidance cues by showing that MDA-MB-231 cells collectively migrate toward the direction of increasing fiber alignment. We anticipate that our easy-to-use experimental platform will enable new studies that explore the effects of graded fiber alignment on cell organization and directed migration in various tissue contexts.

## 2. Results and Discussion

### 2.1 SHG imaging of biopsy samples reveal sub-millimeter gradients in fiber alignment

Over the past two decades, it has been established that cells can migrate along aligned collagen fibers within the tumor microenvironment (TME) in a process called contact guidance[37,38]. To study these responses under controlled laboratory conditions, several teams have developed methods to create uniformly aligned 3D collagen gels and quantify migration metrics, including speed and directionality[36,39]. However, intravital imaging studies have revealed that tissue microenvironments contain local variations in alignment rather than a homogeneous alignment landscape [40]. Therefore, to build physiologically representative contact guidance features, our first objective was to use second harmonic generation (SHG) imaging to quantify the degree of heterogeneity in banked, de-identified triple negative breast cancer (TNBC) biopsy samples. **Figure 1A** shows a representative SHG image of the tumor stroma, highlighting the variation in alignment observed across the 600 μm section. To assess the degree of fiber alignment, the image was divided into three equal regions, and the coefficient of alignment (CoA) was calculated for each region. **Figure 1B** shows that average CoA values ranged from 0.67 ± 0.04 to 0.34 ± 0.07 across the section (slope = 0.02 CoA units/10 μm), with statistically significant differences observed between each section (n=7 samples).

**Figure 1.**
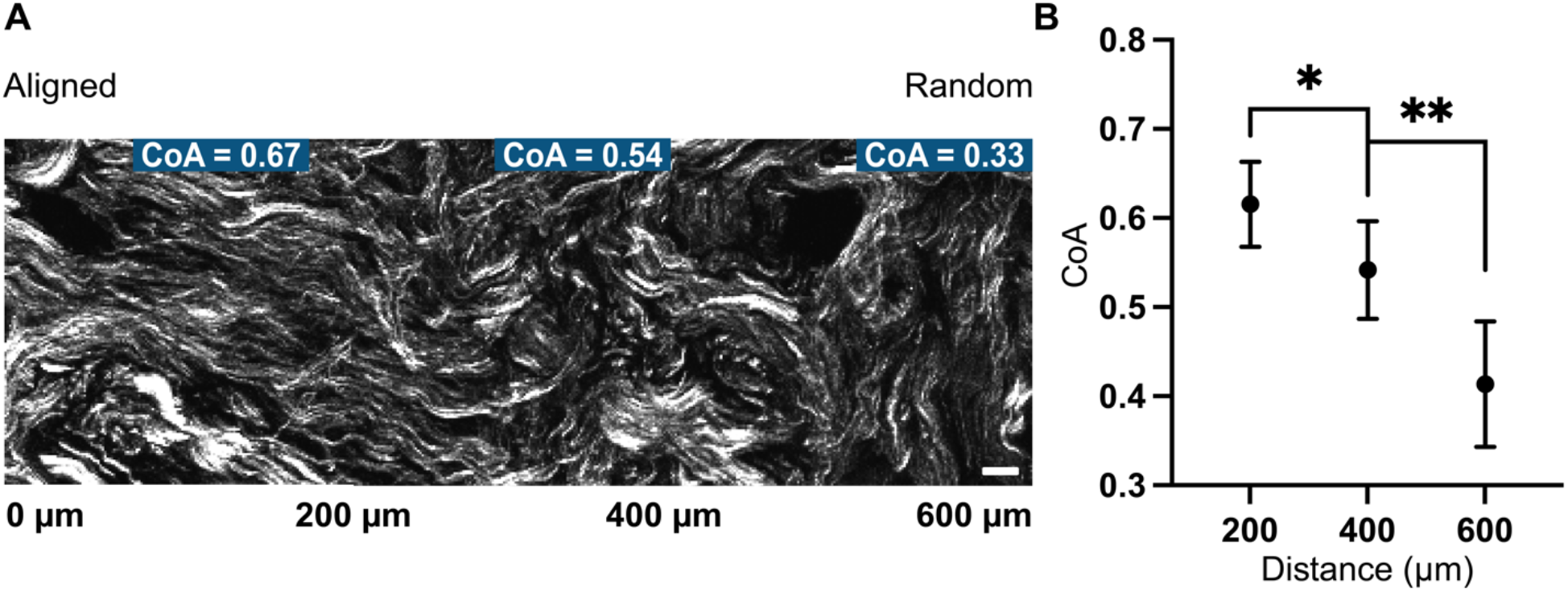
Quantification of fiber alignment gradients in the tumor microenvironment. **A)** A representative SHG image of tumor collagen from a TNBC patient biopsy sample reveals sub-millimeter gradations in alignment, with CoA ranging from 0.67 – 0.33 across the 600 μm section. Scale bar = 25 μm. **B)** CoA is plotted as a function of position along the biopsy sample with statistically significant differences between each section. The data are reported as mean ± standard deviation. *p < 0.05, **p < 0.002 (n=7 samples).

These findings were significant because spatially graded biochemical and biophysical signals present different levels of stimuli across the length of the cell and are known to play an essential role in cell migration [41–45]. These graded cues lead to biased engagement of the signal transduction machinery resulting in asymmetric force generation in the cytoskeleton that promotes directional migration[32]. Hence, the presence of subtle sub-millimeter scale gradients in the biopsy samples prompted us to ask whether alignment gradients could serve as a signal that influenced directional migration.

While alignment gradients resembling the interface between tendon and bone have been achieved using hard synthetic materials like PCL and PLGA through electrospinning, creating fiber alignment gradients in soft collagen-based biomaterials remains a challenge. Cell-mediated traction forces can be used to locally rearrange fiber architecture (see supplemental **Figure S1**); however, the resulting gradient characteristics, including slope and range, are difficult to control. To the best of our knowledge, no existing technologies are capable of microengineering tunable and reproducible alignment gradients in soft collagen biomaterials. Developing a platform with robust biofabrication capabilities is essential to investigate the role of alignment gradients as a guidance cue in the TME and other tissue microenvironments.

### 2.2 Fiber alignment can be controlled by introducing local extensional flows in a microfluidic channel

To address the need for biofabrication techniques to create physiological fiber alignment gradients, we took inspiration from our previous work, where we used local extensional flows to create 3D collagen hydrogels with uniform fiber alignment that persists over 5 mm long distances[46]. This technique involved injecting a neutralized collagen precursor solution into a channel that featured sequential decreases in width, as shown in **Figure 2A**. At the beginning of each segment, the change in geometry introduced a local increase in velocity along the flow direction (quantified by a positive extensional strain rate,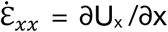, and labeled in blue), which transitioned to a constant velocity state within each parallel-walled segment (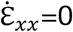 labeled in gray). Here, U_x_ refers to the x component of the velocity vector **U**, and x is the axial direction. In this design, the collagen precursor solution experienced successively increasing extensional strain rate at the entrance of each segment along the flow direction. We discovered that local extensional flows produced highly aligned fibers in the later segments, and the degree of alignment could be controlled as a function of flow rate **Figure S3**. The proposed mechanism was consistent with the behavior of other polymer systems and suggested that a pattern of sequentially increasing extensional flow in the channel induced directional assembly by promoting electrostatic and hydrophobic interactions in the collagen solution that supported globally aligned fiber formation in each segment[15,20,47,48].

**Figure 2.**
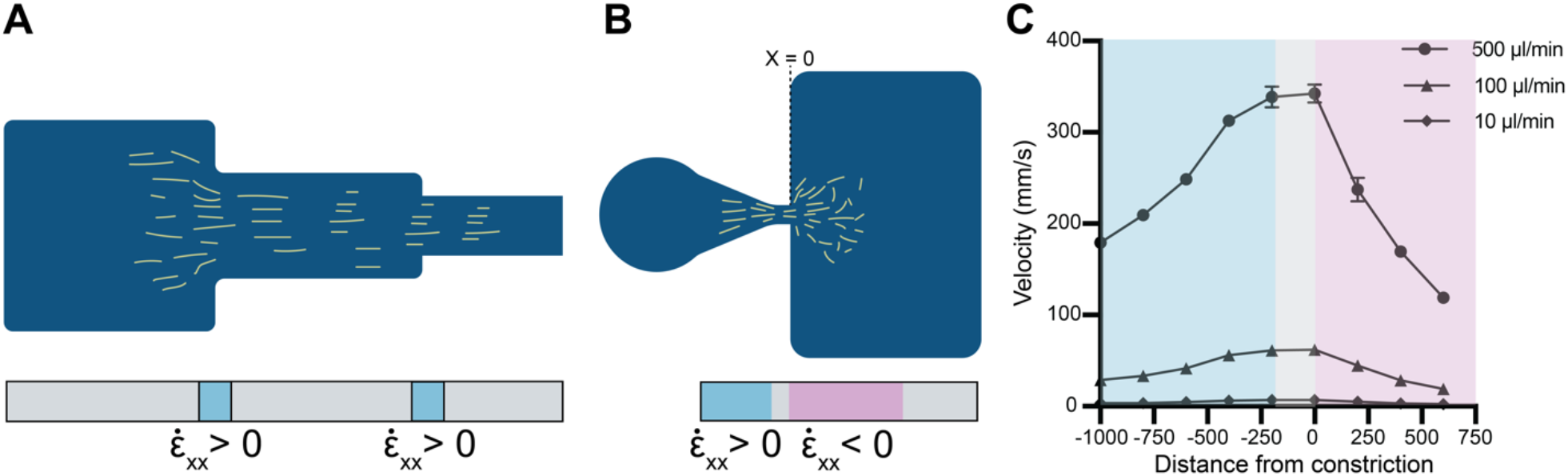
Different extensional flow patterns are introduced by varying the channel geometry. **A)** A positive and zero extensional flow pattern can be achieved using a sequentially constricting design. Light blue regions identify areas of transient stretch due to the velocity increase along the flow direction (+x), while grey areas identify regions with constant velocity. **B)** A pattern of positive, zero, and negative (pink) extensional flows can be achieved in an expanding and constricting channel design. Self-assembling fibers are shown in yellow. **(C)** μPIV data confirms the expected extensional flow pattern for the geometry shown in panel **B** at flow rates Q = 500, 100, and 10 μL min^-1^. Schematics are not drawn to scale.

Based on these findings, we hypothesized that a channel design that incorporated a combination of positive, zero, and negative extensional flows (i.e., positive and negative values of 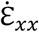 along the flow path) could establish a graded fiber alignment architecture, as illustrated schematically in **Figure 2B**. To confirm that positive (blue) and negative extensional flows (pink) could be generated within a single channel, we designed a geometry with i) a funnel-like constriction, ii) a constant width region, and iii) an expansion well. As shown in **Figure 2C**, we used micro-particle imaging velocimetry (μPIV) to measure the velocity of the neutralized collagen precursor solution in each region with the input flow rates Q = 500, 100, and 10 μL min^-1^. As expected, the extensional strain rate was positive (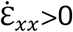, positive extensional flow) in the constriction, zero in the constant width region 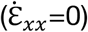, and negative (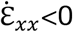, negative extensional flow) upon entering the expansion region. These validation experiments confirmed that the extensional flow patterns the collagen precursor solution experienced along the channel could be controlled as a function of geometry and flow rate.

Next, we tested our hypothesis that a pattern of positive, zero, and negative extensional flows would create continuous gradients in fiber alignment. As shown in **Figure 3A**, neutralized collagen solutions were injected along the path a-a’ at the three flow rates characterized in **Figure 2C**. After gel formation, the resulting fiber microarchitecture was imaged using confocal reflectance microscopy (CRM). We then plotted CoA as a function of position for all flow rates, with the beginning of expansion defined as x=0, and positive x values indicating advancing position into the expansion. As shown in **Figure 3B-C**, fibers were highly aligned (CoA > 0.7) for all flow rates in the parallel-walled region immediately before x=0. The fiber alignment decayed continuously in the expansion for all flow rates but with key differences in the rate of decrease in CoA and the distance over which fibers remained aligned. The green shaded region indicates the distance at which CoA < 0.5 and the fibers were visually unaligned. For Q = 500 μL min^-1^, CoA decreased sharply in the expansion region, transitioning from aligned to unaligned within 500 μm, and created a sub-millimeter scale gradient similar to the TNBC tumor samples. At Q = 100 μL min^-1^, fiber alignment extended 1.2 mm into the expansion, and at Q = 10 μL min^-1^, alignment extended 1.5 mm into the expansion region (**Figure 3B**). **Figure 3C** shows collagen fibers in the pre-expansion funnel region and montages of the corresponding fiber landscapes for each input flow rate.

**Figure 3.**
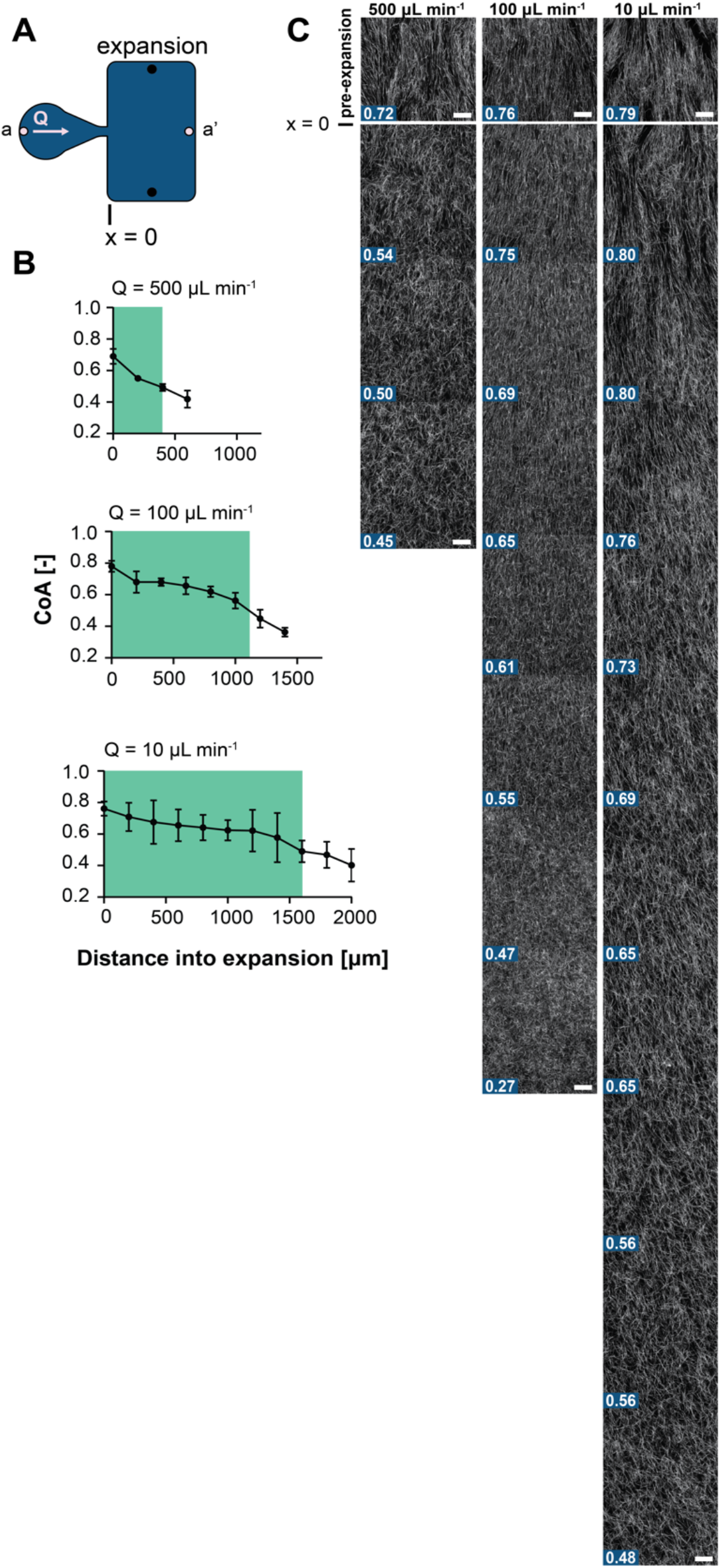
CoA and resulting gradients in alignment as a function of flow rate. **A)** A schematic representation of the channel design is shown with the entrance to the expansion well at x = 0 and defines the flow path a-a’. **B)** The relationship between CoA vs distance into the well is shown for input flow rates Q = 500, 100, and 10 μL min^-1^, with the entrance to the well defined as x = 0. The shaded green boxes mark the distance into the well over which fibers remain aligned, CoA > 0.5. Note that the input flow rate changes the extension of gradient. Results are shown as mean ± s.d. **C)** Representative CRM images corresponding to the data in panel **B** that show continuous CoA gradients after gel formation at each flow rate. CoA values are shown in the bottom left of each image. Scale bar = 25 μm.

Based on these results, we concluded that a pattern of positive, zero, and negative extensional flows in the channel produced defined sub-millimeter to millimeter scale alignment gradients that could be tuned as a function of Q. The mechanism of alignment could be partially explained as follows: in the constricting section, self-assembling collagen subunits were forced into close proximity, stretched, and oriented under the positive extensional flow. As the solution passed into the uniform width region, the velocity was constant (zero extension), and the channel walls constrained the lateral motion of the self-assembling subunits and preserved their orientation. When the collagen solution entered the expansion well, the dramatic increase in the cross-sectional area produced a non-linear decrease in fluid velocity (see **Figure 2C**). The unconstrained flow in the expansion permitted the oriented sub-units to diverge from their upstream positions, followed by diffusive spreading that produced a graded fiber alignment landscape after gel formation [49,50].

One interesting finding was that the highest input flow rate produced the steepest alignment gradient that transitioned from aligned to unaligned within 500 μm and was similar in scale to the sub-millimeter gradients found in TNBC tumor samples (**Figure 1A**). In contrast, the slower flow rates exhibited more gradual changes in alignment, with a longer distance over which the fibers remained aligned. One possible explanation for this behavior could be described by considering the Deborah number (De), which is a ratio between the solution relaxation time (*λ*) and the flow time scale (*τ*), De = *λ*/*τ*. When De > 1, the elastic behavior of the polymer solution dominates, while viscous, liquid-like behavior dominates as De << 0 [51]. At a flow rate of 500 μL min^-1^, De = 4.1, suggesting that the elastic response of the collagen solution in the expansion could contribute to the sharp gradient in alignment. Conversely, at the lower flow rates tested (Q = 100 and 10 μL min^-1^) the response of the solution was dominated by viscous rather than elastic effects (De = 0.74 and 0.08, respectively), producing a gradient with a less pronounced slope. Although the mechanism of gradient formation remains an active area of investigation in our laboratory, the primary goal of this work was to highlight the concept that simple variations in channel geometry can create sub-millimeter to millimeter scale gradients in fiber alignment and provide unique biofabrication capabilities in soft materials.

To expand the material options available with our approach, we also show that our microengineering technique is not limited to collagen gels. We have successfully used collagen/fibronectin, collagen/hyaluronic acid (HA), and glycated collagen to provide a range of options to engineer in vitro models to better represent the architecture and composition of the target tissues (**Figure S4**). Our technique is also compatible with photoactivatable methacrylated collagen that allows the matrix modulus to be tuned to match target tissue through exposure to UV light (see **Figure S5**). Notably, the collagen microarchitecture is maintained before and after UV exposure, as shown in **Figure S6**. Our approach is also compatible with published methods that control fiber diameter by sequentially varying the gelation temperature from room temperature to 37°C [52–54]. Thus, our versatile microengineering technique provides a wide range of options to achieve in vitro models that faithfully replicate the architecture and composition of natural tissues. We anticipate that this technique can be applied to various fibrous ECM materials, including fibrin and elastin, to further support advancements in soft material biofabrication.

### 2.3 Matrix fiber density and pore diameter are not affected by collagen CoA

Gradations in matrix pore diameter and fiber density are well-studied taxis cues that guide cell migration through the ECM[55–57]. Thus, after establishing a method to create 3D collagen gels with tunable CoA gradients, we investigated whether CoA influenced pore size and fiber density. We first categorized fiber architecture into three groups: low (0.25 < CoA < 0.5), medium (0.5 ≤ CoA < 0.7), and high (0.7 ≤ CoA < 1). To calculate the pore diameter in each group, we used a standard image processing algorithm to allow for benchmarking against other work. **Figure 4A** shows no statistical differences in pore diameter between the low, medium, and high CoA categories, and the average pore across the groups was 2.72 ± 0.15 μm. Next, we investigated whether CoA influenced fiber density, as measured by the fiber fraction. The relationship between CoA and fiber fraction is shown in **Figure 4B**, and no statistical differences were found among the three alignment categories, with an average fiber fraction of 0.17 ± 0.02. These values are consistent with those reported in the literature using similar collagen concentrations and gelation conditions [58,59]. Taken together, these analyses provide evidence that fiber alignment can be controlled without influencing bulk pore size or fiber density using our flow-based biofabrication approach.

**Figure 4.**
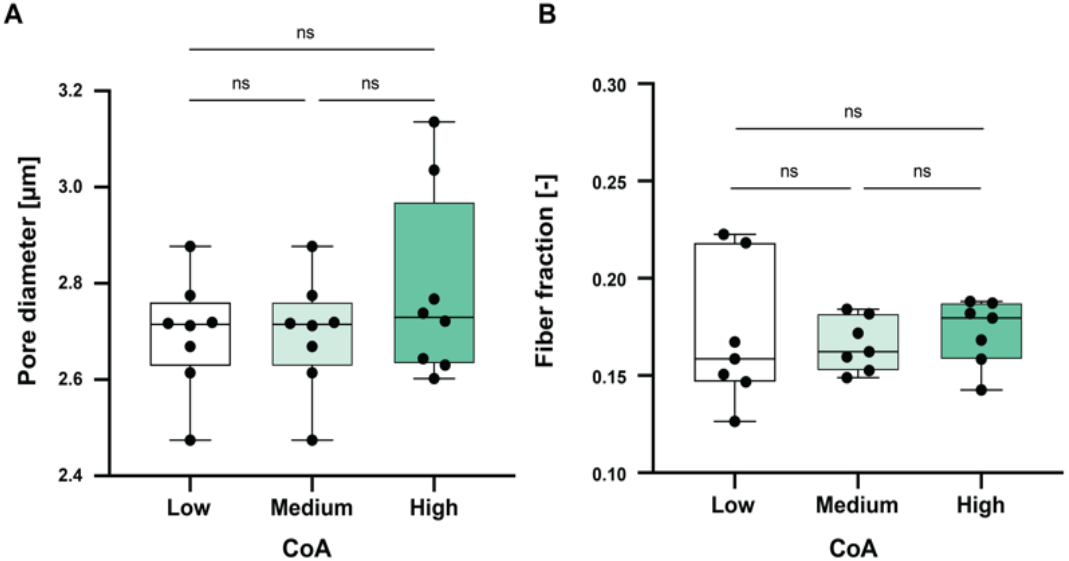
Matrix pore diameter and fiber fraction as a function of CoA. Collagen CRM images were divided into 3 categories, high (green, 0.7 < CoA ≤ 1), medium (light green, 0.5 ≤ CoA ≤ 0.7), and low (white, 0.25 ≤ CoA < 0.5) alignment. **A)** The box and whisker plots show the pore diameters in the three CoA categories and ANOVA analysis revealed no statistical differences between groups. **B)** The box and whisker plots show fiber densities in the three groups and ANOVA analysis concluded the values are not statistically different. Box and whisker plots show median, 1^st^ and 3^rd^ quartiles, and the minimum and maximum values. n = 7 independent gels per group. ns = not statistically significant.

These findings are relevant because ligand density is directly related to the fiber density in the matrix, and our data show that the fiber density (measured by fiber fraction) is independent of CoA [53,60]. Fiber density and porosity are often used as a proxy for the matrix storage modulus [61–63], and our results indicate that these properties are independent of CoA, suggesting that bulk mechanical properties may not be linked to CoA in our system. Taken together, our system could be used to quantify how cell migration is influenced by fiber alignment gradients independent of underlying differences in bulk porosity or density that could confound experimental findings.

### 2.4 A reversibly sealed microchannel allows direct access to microengineered 3D collagen gels

After quantifying relationships between CoA, porosity, and fiber density in our gels, we sought to quantify how cells respond to fiber alignment gradients in our microfluidic platform. Microfluidic approaches are well-recognized as experimental tools that enable precise control over the cellular microenvironment[64–66]. Conventional microfluidic systems are permanently sealed, and the introduction of cells requires that they are included in the self-assembling gel solution or added through ancillary seeding channels are used[67–70]. Incorporating cells in the gel solution limits the parameter space (e.g., gelation temperature, ionic strength, pH, mechanical strain, or photoinitiator concentration) that can be used to control matrix properties. Incorporating additional seeding channels can complicate designs and introduce sources for bubble formation or clogging that can decrease experimental yield [71,72]. Our microfluidic design simplifies the process of adding cells to the gels by using a BSA passivated PDMS lid that is sealed reversibly against a through-cut PDMS sheet that contains the desired channel geometry. The completed channel is comprised of a glass coverslip on the bottom, PDMS side walls, and a PDMS lid containing cored access ports. The PDMS lid is peeled off after gel formation, providing full access to the microengineered collagen matrix[20]. Using CRM, we verified that the fiber microarchitecture was not disrupted by comparing fiber alignment before and after the peel off process. **Figure S7** shows the change in CoA, ΔCoA = 0.02 ± 0.03, with a maximum change of 0.06 CoA.

Our previous work shows that the open access design also allows specialized modules to be attached to the gel layer. These include permeable cell culture membranes, flow modules, and gel deposition modules to support layer-by-layer assembly [46]. Here, we add cells directly to the gel or use a second PDMS sheet containing a patterning aperture sealed against the PDMS channel layer to define the location of the added cells. This technique is also compatible with 3D printed or automated dispensing methods. Thus, our approach combines the advantages of conventional microfluidic systems but simplifies the operations typically associated with these approaches.

### 2.5 HUVECs and MDA-MB-231 cells migrate directionally on gradient gels

After confirming that the lid could be peeled off with a minimal change in CoA, we seeded HUVECs directly on random and gradient gels to verify that the single cells could sense and respond to fiber architecture (see **Figure 5A)**. Cells closer to the entrance of the expansion well (x=0) experienced CoA values close to 0.7, while the rear of the expansion region well was less than 0.5. **Figure 5B** shows HUVECs cell tracks on an unaligned control gel and a microengineered gradient gel over 8 hours. For each group, we calculated directionality (D), defined as the ratio between the shortest distance (L) between cell position at the beginning and end of the experiment compared to the total path length traveled (d), D = L d^-1^. This metric can range from 0 (low) to 1 (high).

**Figure 5.**
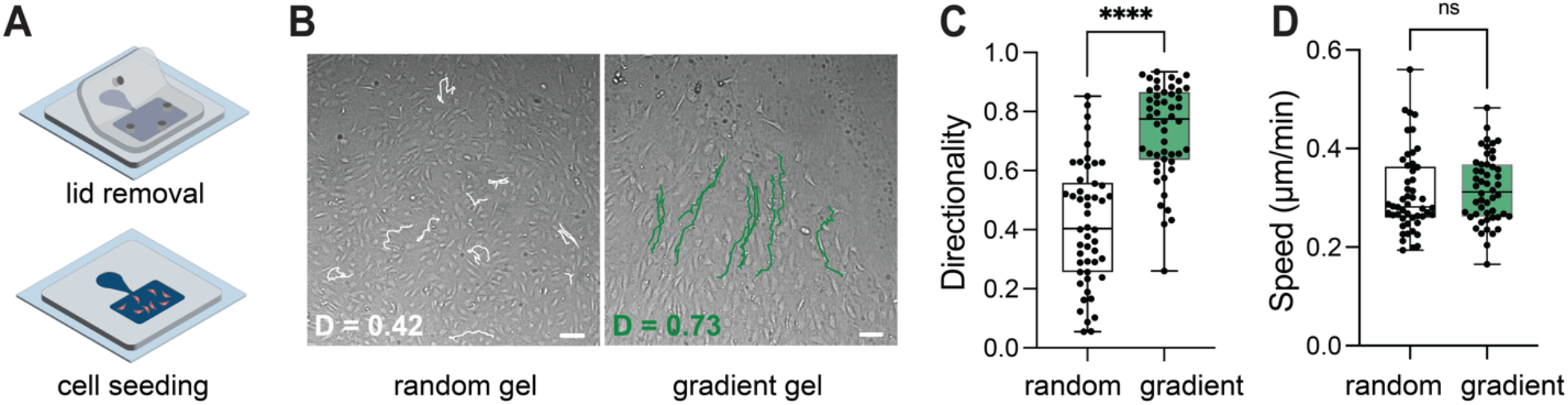
Migration characteristics of HUVECs on gradient and unaligned collagen gel. **A)** Schematic representation of detaching the PDMS lid to access the engineering hydrogel and seeding cells in the well region. Note, the gel is confined to the footprint of channel, and the height is defined by the thickness of the sheet. **B)** Migration tracks of ten randomly selected cells on an unaligned gel are shown in white and gradient gel shown in green. The cells on unaligned gel exhibit a directionality of 0.42, representing no preferential directionality while the cells on gradient gel have a directionality of 0.78 indicating preferential movement. Scale bar = 25 μm. **C)** Box and whisker plots show the directionality of 50 cells on random and gradient gels. Statistical comparison of the data show cells on gradient channel exhibit higher directionality than cells on unaligned collagen gels. **D)** Box and whisker plots show the speed of 50 cells on random and gradient channel, with no statistical differences between the groups. ****p < 0.0001, ns = not statistically significant, n= 50 cells.

The cells on the gradient gel exhibited directional migration toward increasing alignment with D = 0.73 ± 0.16, while the cells on unaligned collagen gels exhibited D = 0.42 ± 0.21. The average speed was not statistically different between the aligned (0.31 ± 0.08 μm min^-1^) and random gel conditions (0.32 ± 0.06 μm min^-1^), but time-lapse imaging shows that cells on the gradient gel (Video S1) migrated directionally with limited backward motion, while cells on the unaligned gel exhibited random migration (Video S2). This behavior is similar to other contact guidance studies using cancer cells where directionally increased in aligned gels, but speed was consistent [73]. Although there were alignment gradients present in the system, we reported pooled, population-level migration metrics rather than region-specific motility metrics because the purpose of this experiment was to demonstrate that single cells seeded on the gels could respond to the fiber architecture created in our system and set the stage for future studies.

After showing how cells responded to alignment in our system, our next goal was to test the hypothesis that fiber alignment gradients could direct cell motility similar to classical taxis cues. To do so, we removed the PDMS lid after gradient gel formation and replaced it with a PDMS sheet that contained a circular patterning aperture [74]. An MDA-MB-231 cell aggregate was added to the aperture and settled onto the gel. **Figure 6A** shows a representative image with fluorescently labeled MDA-MB-231 positioned within the footprint of the aperture. Cells at the left edge of the aperture experienced a CoA of > 0.6 while the CoA < 0.5 (random) at the right edge. We observed the collective movement of the cell aggregate over five days, capturing images once every 24 hours and measuring the positions of the left (l_n_) and right (r_n_) edges of the cell aggregate. We identified an enclosed cell area for each time point using image processing. Although this analysis looks at an enclosed area, cells do migrate into the gel. To assess the relative cell migration in each direction, we calculated a metric called Ω_n_, which compared the ratio of Δl_n_ (the change in the left edge position over 24 hours) to Δr_n_ (the change in the right edge position over 24 hours). l_24_ and r_24_ positions were used as the origin to eliminate any initial bias if the aggregate was not centered in the aperture after seeding. As shown in **Figure 6B**, Ω_n_ > 1 indicates biased collective movement toward increasing CoA. As shown in **Figure 6C**, Ω_48_ = 1.1, indicating a slight leftward bias, and 24 hours later, Ω_72_ = 1.6, indicating continued leftward progression of the cell sheet. The highest Ω_120_ = 3.2 was measured at 120 hours after seeding and represented a distinct difference in directional cell migration toward increasing alignment. We observed that cells appeared to move along the fiber direction with minimal migration orthogonal to the fibers. Based on these preliminary findings, we conclude that the gradients in fiber alignment developed in our system warrant further exploration as in contact guidance.

**Figure 6.**
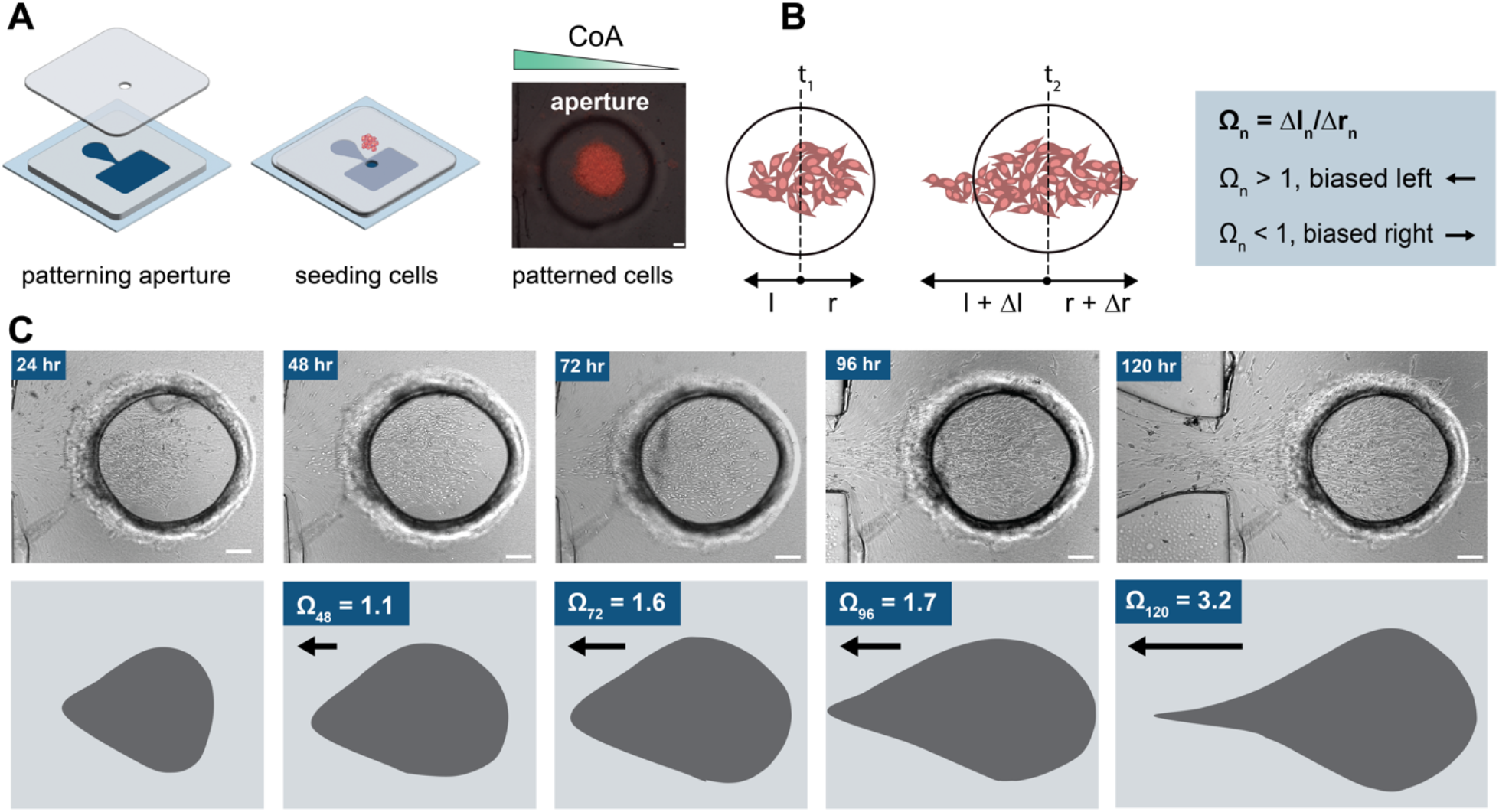
MDA MB 231 cell patterning and biased migration on graded collagen gels. **A)** A schematic illustrating the process of patterning cell aggregates through the aperture with a micrograph of fluorescently labeled cells patterned on a collagen gel. **B)** The schematic describes how cell movement was tracked by marking the left and right ends of the advancing cell front, with the center of the window considered the initial point. **C)** Time-lapse micrographs taken every 24 hours, displaying the progression of the enclosed cell area, highlighted by a blue envelope. Cells show a marked tendency to move towards areas with increasing fiber alignment, characterized by Ω_n_, a ratio representing the advancement in enclosed cell area advancement from left to right. Scale bar = 200 μm

Although contact guidance along topographic features has been observed in several tissue microenvironments, the mechanism that promotes directional migration when no intrinsic asymmetry is present (e.g., no preferred directionality along the aligned fibers) is not entirely understood. One explanation is through the anisotropic mechanical resistance hypothesis that explores the migration of dermal fibroblasts in fibrin gels containing uniform fiber alignment with constant adhesion strength and porosity[75]. Crosslinked and stiffened gels exhibited anisotropy in the mechanical resistance along and perpendicular to the fiber direction that could be sensed by the cells. Although these results were generated in aligned fibrin gels, there are direct analogs to crosslinked, aligned collagen gels. Another possible explanation, based on experimental findings in MDA-MB-231 cells and supported by computation modeling, is that the degree of collagen fiber alignment directly influenced the elongated versus rounded cell morphology and dictated their migration mode (mesenchymal or ameboid). Cells in contact with highly aligned fibers exhibited a high aspect ratio and mesenchymal migration, while cells interacting with unaligned fibers appeared rounded and exhibited amoeboid-like migration. Cells on moderately aligned collagen switched between mesenchymal or ameboid modes. The elongated cells appeared more sensitive to contact guidance, while low aspect ratio cells largely ignored ECM cues[76]. Both studies aimed to identify the sensing mechanism in contact guidance and were performed in uniformly aligned hydrogel environments.

Our experimental platform offers a complementary approach to investigate contact guidance mechanisms and cell responses by providing continuous, spatially graded fiber alignment landscapes found in vivo but difficult to recreate in soft materials. Here, cell populations are exposed to different levels of alignment (rather than uniform alignment) and are analogous in presentation to established cell taxis cues. Thus, this platform allows new questions to be explored. An additional strength of our platform lies in the simplicity of the geometry-based biofabrication approach, which does not require specialized equipment such as a bioprinter or electrospinning setup. The microfluidic device can be made using a laser cutter and PDMS molding, and the basic constricting and expanding channel design provides a straightforward and accessible approach to achieve fiber alignment gradients. The main goal of this work was to introduce the platform and demonstrate that gradient characteristics can be tuned as a function of flow rate in an expanding and contracting microfluidic channel. Future development work will focus on establishing geometry and flow rate based design rules to create more complex fiber architectures further advance biofabrication capabilities.

## Conclusions

This work introduces a simple geometry-based microfluidic approach to biofabricate programmable sub-millimeter to millimeter scale fiber alignment gradients within a 3D collagen-based material library. Our reversibly sealed channel design allows direct access to the microengineered gels and enables controlled cell patterning at defined locations, facilitating directed migration studies in graded fiber alignment landscapes. As a demonstration of the platform, we provide evidence that patterned MDA-MB-231 cell aggregates respond to subtle gradients in fiber alignment by collectively migrating toward the direction of increasing fiber alignment. We anticipate that our first-of-its-kind biofabrication technique to engineer alignment gradients will enable new studies to explore how cells interpret and respond to heterogeneous biophysical environments.

## Supporting information

HUVECs exhibit a random walk pattern during migration on an unaligned collagen gel

Directional migration of HUVECs on collagen fiber alignment gradient

## Acknowledgements

The authors acknowledge Thomas Gaborski for assistance with cell motility analysis, Steven Day for μPIV characterization, Vincent Mei and Christopher Lewis for assistance with rheometry and UV exposure studies. The authors also thank Meng-Chun Hsu, Ann Byerley, Madeleine Goulet, Priya Benny Chiriyankandath, Aidan Hughes, Lauren Audi, and Anna Carter of the Biological Microsystems Laboratory at RIT for insightful discussions. This work was partially supported by the NIH under grant numbers R21GM143658, R16GM146687, and NSF grants CBET 2150798 and 215099. This content is solely the responsibility of the authors and does not necessarily represent the official views of the National Institutes of Health or the National Science Foundation.

## Materials and Methods

### Banked patient samples and second harmonic generation imaging

We imaged tumor-stroma interface in biopsy samples of triple-negative breast cancer (TNBC) patients with Second Harmonic Generation (SHG) imaging. These patients were identified from the pathology files at the University of Rochester Medical Center (URMC), spanning 2009-2020. The use of patient samples was approved by the Institutional Review Board at the University of Rochester (IRB RSRB00069270).

To image the collagen fibers in the TME, we used a Spectra-physics MaiTai Ti:Sapphire laser directed through an Olympus Fluoview FV300 scanner and a BX61WI upright microscope. The laser’s circular polarization was confirmed to ensure equal excitation of all fiber orientations. We focused the laser through an Olympus UMPLFL20XW water-immersion lens, which also collected the backward-scattered SHG signal. This signal was separated from the excitation beam, filtered, and detected by a photomultiplier tube. The forward-scattered SHG signal was collected by an Olympus 0.9 NA condenser, reflected, filtered, and detected by another photomultiplier tube. We collected 512×512 pixel images as a Z-stack in 3 μm steps and then maximum-intensity projected to form a single forwards-detected image. The fiber alignment in a forward detected image was assessed by dividing the image into 200 × 200 μm regions and calculating the coefficient of alignment (CoA) for each. CoA values can range from 0.25 – 1, representing low to high fiber alignment, with a value of 0.5 serving as the threshold for alignment

### Resealable microchannel fabrication

We first created a mold using standard soft lithography techniques[20,46,77]. Briefly, a negative photoresist, SU-8 3050 (Kayaku Advanced Materials, Massachusetts, USA), was spin-coated on a 100 mm diameter silicon wafer (University Wafers, South Boston, USA) to a thickness of 200 μm. The spin-coated height (200 μm) defined the height of the microchannel. The photoresist was then soft-baked at 95 °C for 60 mins, followed by exposure to 365nm UV (250mJ/cm2) through a high-resolution printed photomask. The wafer was baked at 95 °C for 5 mins, cooled to room temperature, and immersed in a SU-8 developer solution to remove the non-crosslinked photoresist. The wafer was then rinsed with isopropanol and dried under pressurized air.

We then mixed the PDMS base and crosslinker (Sylgard 184, Dow Corning, Midland USA) in a 10:1 ratio and degassed the mixture in a vacuum chamber for 30 minutes to remove bubbles. We poured the degassed PDMS on the silicone mold and created a stack consisting of a transparency sheet, metal plate, and a 6 kg mass. The mass pushed excess PDMS away from the SU-8 features and created a through-cut channel geometry, with a height of the feature defined by the SU-8. The stack was cured at 80°C for 1 hour on a hotplate. After cooling to room temperature, the transparency sheet was carefully removed from the mold. The through-cut PDMS sheet was then peeled from the mold and inspected. The PDMS was cut into individual sections and cleaned with 70% ethanol solution and DI water. The through-cut PDMS sheet was treated with air plasma on one side and bonded to a glass coverslip to form a microchannel with 200 μm height. A PDMS lid with ports for injecting collagen was sealed against the PDMS channel footprint using conformal contact.

To fabricate PDMS lids, we laser cut lid shapes into 1.5 mm thick acrylic sheets and attached those to a silicon wafer using the PSA film, yielding six square-shaped cavities, each having 20 mm sides and a 1.5 mm depth. Following this, we filled the silicon mold with degassed PDMS, allowing us to create lids of 1.5 mm thickness. We then heated the silicon mold 80°C for 1 hour and then allowing it to cool to room temperature. To finish, we removed the lids from the mold and cleaned them using a 70% ethanol solution. The clean lids were then stored with protective tape applied to both faces.

### Coverslip functionalization and PDMS passivation

To enhance the attachment of collagen to the coverslip, we functionalized coverslips with poly(octadecene maleic alt-1-anhydride) (POMA) using established protocols [20,78,79]. Briefly, the coverslips were cleaned with 70% ethanol and dried under a pressurized air stream. The coverslips were then exposed to O_2_ plasma for one minute at 600 mTorr (Harrick Plasma, NY, USA) to activate their surface. Next, the coverslips were immersed in a 2% v/v solution of aminopropyltriethoxysilane (APTES) (Millipore Sigma, VT, USA) and placed on a rocker for 5 minutes. Following this step, the coverslips were rinsed with ethanol, dried under pressurized air, and baked on a hot plate for 10 minutes at 110°C. The POMA solution was prepared by dissolving poly(octadecene maleic alt-1-anhydride) (Millipore Sigma, Burlington, USA) in 99.9% tetrahydrofuran (Millipore Sigma, Burlington, USA), and the coverslips were spin-coated with this solution. Finally, the POMA-coated coverslips were placed on a hot plate for 1 hour at 120°C.

To prevent the unwanted attachment of collagen to the PDMS lid, the surfaces were passivated by immersing them in 4% w/v Bovine Serum Albumin (BSA) in 1x PBS for 4 hours at 4°C. The PDMS lids were rinsed 3X with PBS, allowed to air dry, and stored in a closed Petri dish until use.

### Collagen solution preparation and injection

Atelo collagen: A total of 416 μL of bovine atelo Type I collagen solution (Neutragen, Advanced Biomatrix, CA, USA) was diluted with 429 μL of ultra-pure water, 55 μL of 0.1M Sodium hydroxide (VWR, PA, USA), and 100 μL of 10X phosphate-buffered solution (ThermoFisher Scientific, MA, USA) to achieve a final collagen concentration of 2.5 mg/ml. The pH of the neutralized collagen solution was adjusted to 8.7-9.2 using 0.1M NaOH and measured with a pH probe (A511 Orion Star, ThermoFisher, USA) after each preparation.

Methacrylated collagen: Photoactive methacrylated Type I Bovine Telo collagen kit was purchased from Advanced Biomatrix (Neutragen, Advanced Biomatrix, CA, USA). A 3 mg/ml collagen stock solution was mixed with the neutralizing solution and the photoinitiator Irgacure 2959 to achieved a final collagen concentration of 2.7 mg/ml following the recommended protocol, resulting in a pH between 7.4 and 7.6. After the collagen mixture polymerized, we further crosslinked the collagen hydrogel by exposing the device to 365 nm UV light at 10 mW cm^-2^ for 275 seconds.

A syringe pump (New Era Pump Systems, NY, USA) was used to inject neutralized collagen solution into PDMS microchannels in two phases. During the first phase, collagen was injected into port 1 at a flow rate of 500 nL min^-1^ using a 20-gauge dispensing needle (Grainger Industrial Supply, IL, USA) while all ports were open[80]. Preloading the channel with collagen prevented air bubbles from forming during the next step. In the second phase, ports 2 and 4 were closed, and collagen was injected at varying flow rates (Q = 10 μL min-1, 100 μL min-1, and 500 μL min-1) from port 1 to port 4. The injections were conducted at room temperature (21°C). The chip was then incubated in a petri dish at 37°C for 1 hour to facilitate collagen polymerization. A damp Kim wipe (Kimtech, Kemberly-Clark Professional, TX, USA) was placed in the petri dish to prevent evaporation.

### Microparticle image velocimetry

Microparticle image velocimetry (μPIV) to measure the velocity of the neutralized collagen solution. The measurements were performed using a Nikon Eclipse TE2000-S microscope equipped with a pulsed laser (TSI Incorporated, MN, USA). We injected a mixture of 2.7 mg/mL of neutralized collagen solution and 5 μm fluorescent polystyrene latex particles (0.1% w/v) (Magsphere, CA, USA) at Q = 500, 100 and 10 μL min^-1^ from the inlet of the microfluidic chip using a syringe pump. To obtain the PIV data, we performed measurements on three independent samples in three different regions. We analyzed ten pairs of images, captured at 10 μs intervals within a 700 × 700 μm region centered on the microchannels. The velocity of the neutralized collagen solution was calculated using TSI Insight 4G software (TSI, MN, USA).

### Confocal imaging and analysis of collagen fibers

We used a Leica SP5 laser scanning confocal microscope equipped with a 40X water immersion objective and 1.75X optical zoom to image collagen fibers. We captured 2 μm image stacks consisting of 13 images at z = 30 μm from the coverslip. Collagen fibers were imaged in reflectance mode using a 488 nm laser. To analyze the images, we projected the image slices onto a single plane using the average projection feature in FIJI (NIH, USA) before analysis.

We used an Olympus IX-81 microscope with a 4X objective for cell patterning imaging. To conduct the migration experiment, we used the Leica SP5 microscope with a 20x objective and an incubated stage with temperature and gas control. We loaded the coverslips containing the gradient channels and patterned cells onto a 6-well plate and placed them on the microscope stage. The 6-well plate was incubated on the microscope stage for 1 hour before starting imaging.

We analyzed confocal reflectance microscopy images using LOCI CT-FIRE, a MATLAB-based code [81], to measure fiber alignment. We obtained a histogram of fiber angle distribution in 15° bins for each image and calculated the coefficient of alignment (CoA) as the ratio of the number of fibers within ±15° of the mode on the histogram to the total number of fibers. A CoA > 0.5 indicated aligned fibers, and CoA < 0.5 indicated random fiber arrangement. These methods allowed us to accurately measure fiber alignment and assess cell migration in our experiments.

### Pore diameter, fiber density, and cell motility characterization

In order to analyze the pore characteristics of our gels, we employed a method previously described by Taufalele et al [53]. The initial step involved projecting an image stack onto a single plane. Our images measured 221.64 × 221.64 microns in the x and y directions, with a pixel/μm ratio of 4.623. In our case, each z-stack consisted of 13 slices, which were projected onto a single plane using the average projection in Fiji. The projection was imported to MATLAB, the Erode Cutoff was set to one, and a top-hat filter was applied to adjust contrast. To determine the fiber fraction, the projected image was binarized, and a ratio of bright pixels to total pixels was calculated and reported as the fiber fraction for each group of images corresponding to the high, medium, and low CoA categories.

To track the migration of the cell front, we first cropped the 2024 × 2024 cell aggregate microscope image to a 1734×1024 image that included the cell-containing region, using Fiji. We then aligned all images using a translating function so that the center point of the patterning aperture coincided across all images. This center point was used at the origin to measure the cell distance to the left and right. To identify the cells within the image, we applied a 3×3 second derivative convolution filter, which helped subtract the patterning aperture from the image. Next, we used a variance filter with a 2-pixel radius to enhance the contrast between cells and their surroundings, improving the tracking of the cell front. Finally, we converted the resultant image into a binary black and white frame (enclosed area) using the Fiji Otsu threshold filter, allowing us to calculate the distance of both cell fronts from the previously marked center point.

### Cell culture and cell aggregate formation

MDA-MB-231 cells (ATCC, VA, USA) were cultured in DMEM (11995065, Gibco, ThermoFisher, MA, USA) supplemented with 10% Fetal Bovine Serum (FBS). Human Umbilical Vein Endothelial Cells (HUVECs) (ATCC, VA, USA) were cultured in Endothelial Cell Basal Medium (EBM) supplemented with the Endothelial Growth Media kit (EGM-2) (CC-3162, Lonza, Basel, Switzerland). All cells were maintained at 37°C and 5% CO2 and used between passage number 3-7. Loose MDA-MB-231 cell aggregates were formed by adding 15000 cells into an untreated U-shaped bottom 96-well plate and incubated at 37°C for two days. The resulting cell aggregates were then labeled with CMDA or unlabeled.

### Cell patterning

After collagen gel formation, the functionalized PDMS lids were carefully peeled away, and 500 μL of media was added to the collagen gel, which was then placed in an incubator overnight. Following incubation, the excess liquid was removed from the collagen surface, and a 200 μm thick PDMS sheet containing a laser cut 500 μm diameter patterning aperture was positioned on the collagen, providing direct access to a defined region of the gel.

To pattern a cell aggregate, the aggregate was removed from a non-treated U-shaped bottom plate using a sterile spatula and added to the patterning aperture, settling on the collagen gel within the aperture footprint.

### Cell labeling and immunostaining

CMRA cell tracker dye (ThermoFisher Scientific, MA, USA) was diluted to a concentration of 25 μM in cell media and added to the culture flask to enable live tracking of cells. The cells were incubated with the dye for 30 minutes, followed by replacing the media containing the CMRA stain with a culture medium.

For immunostaining, the cells were fixed with 4% paraformaldehyde (ThermoFisher Scientific, MA, USA) and permeabilized with 0.1% Triton X-100 (J63521, Alfa Aesar, MA, USA) for 15 minutes. The permeabilized cells were then blocked with 4% Bovine Serum Albumin (BSA) (VWR, PA, USA) for 30 minutes, and each step was followed by washing the cells with PBS Tween 20 (ThermoFisher Scientific, MA, USA).

### Statistical analysis

The CoA of collagen alignment data was displayed as the average ± standard deviation from n=3 independent gels. We examined n=7 specimens for SHG tumor imaging to evaluate the in vivo gradient in collagen fiber alignment. To assess fiber fraction, porosity, and CoA differences, we studied eight samples per condition. We first used the Shapiro-Wilk normality test to verify the data has a normal distribution. We then applied a standard one-way ANOVA and a Tukey post hoc test to identify statistically significant differences. Box and whisker plots were presented as median,1st and 3rd quartiles, and min and max values. A p-value less than 0.05 was considered statistically significant. All statistical analyses were performed using GraphPad Prism 9.0 software (GraphPad Software, CA, USA). ns = not significant, *p < 0.05, **p < 0.002, ***p < 0.0002, and ****p < 0.0001.

