## Supplementary information for "Microengineering 3D Collagen Matrices with Tumor-Mimetic Gradients in Fiber Alignment"

### Supporting Information

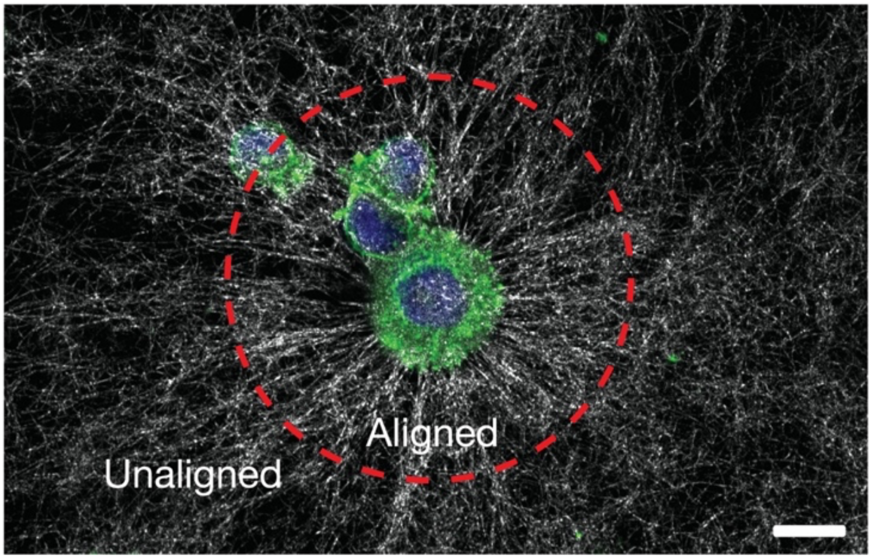


**Figure S1:** Shows the COL1 fibers remodeled by MB 231 cells through traction forces. It is seen that fibers are aligned next to the cell boundary, with fiber alignment decreasing with distance from the cell (Scale bar = 25 μm).

**Figure S2:** Shows the velocity vectors obtained from PIV in the three regions of interest (Constriction, constant width, and expansion). The length of the vectors is representative of the velocity magnitude experienced by the neutralized COL1 solution as it flows through the microfluidic chip.

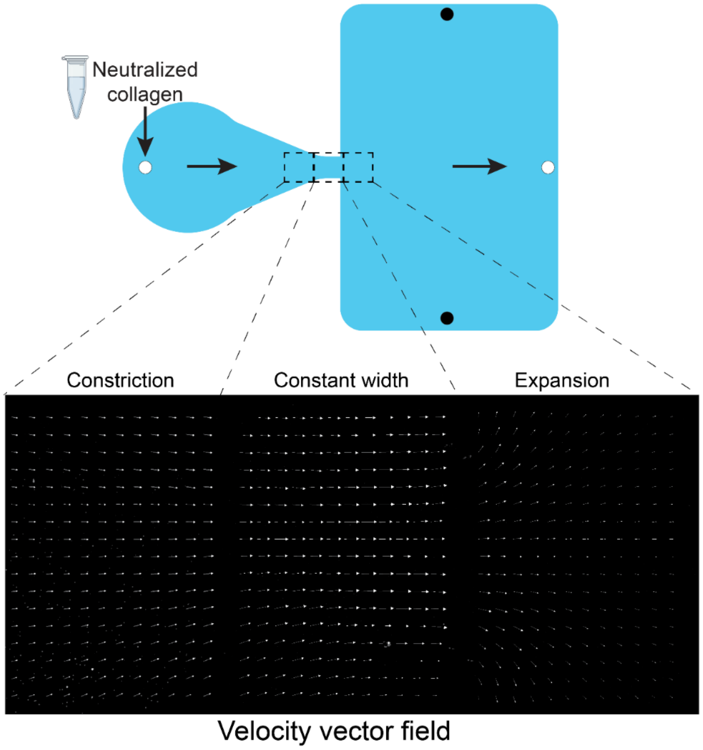

**Figure S3: A)** Schematic of the sequentially constricting channel design. The change in geometry creates a local extensional strain in the collagen 1 prepolymer solution that promotes 5mm long segments of uniform alignment after gelation. **B)** CRM images of collagen in the yellow highlighted boxes show differences in CoA along the channel. Scale bar = 25 μm.

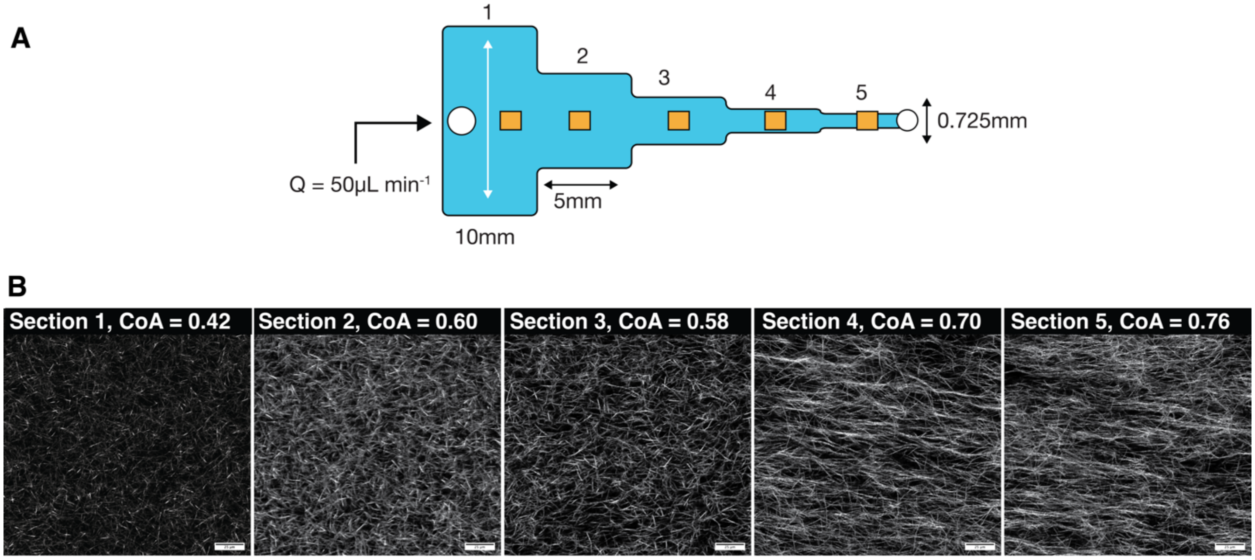


**Figure S4:** Confocal images of aligned collagen gel with HA, Ribose, Fibronectin Rat Tail and collagen incubated at room temperature before polymerization at 37 °C.

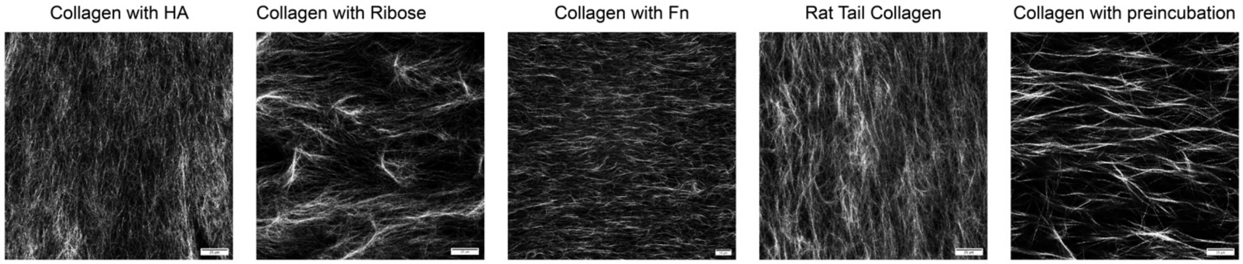

**Figure S5:** Stiffness modulus of collagen with rheology during polymerization and post polymerization cross linking by exposing to 365 nm UV light

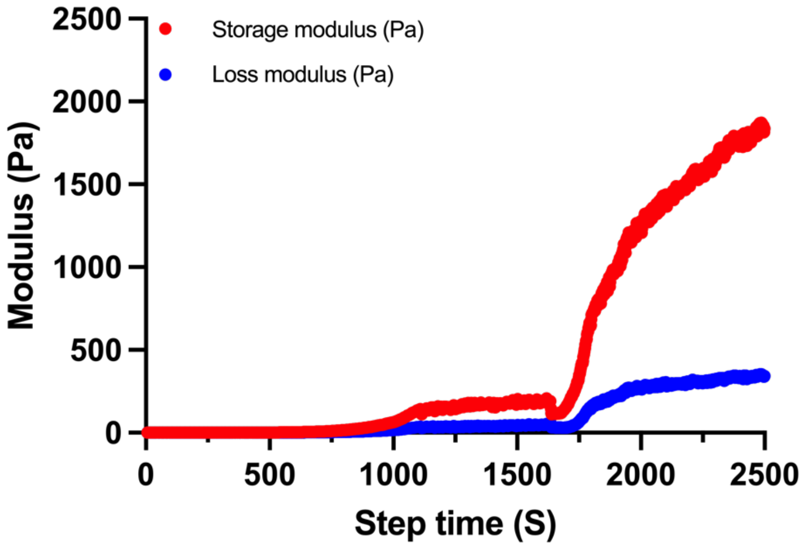


**Figure S6: A)** Figure shows the graph of CoA vs distance from constriction for the ColMA before and after exposure (n=3). **B)** Representative images of COLMA fibers in the constriction (0 µm), 400 µm and 800 µm in the expansion region (Scale bar = 25 µm).

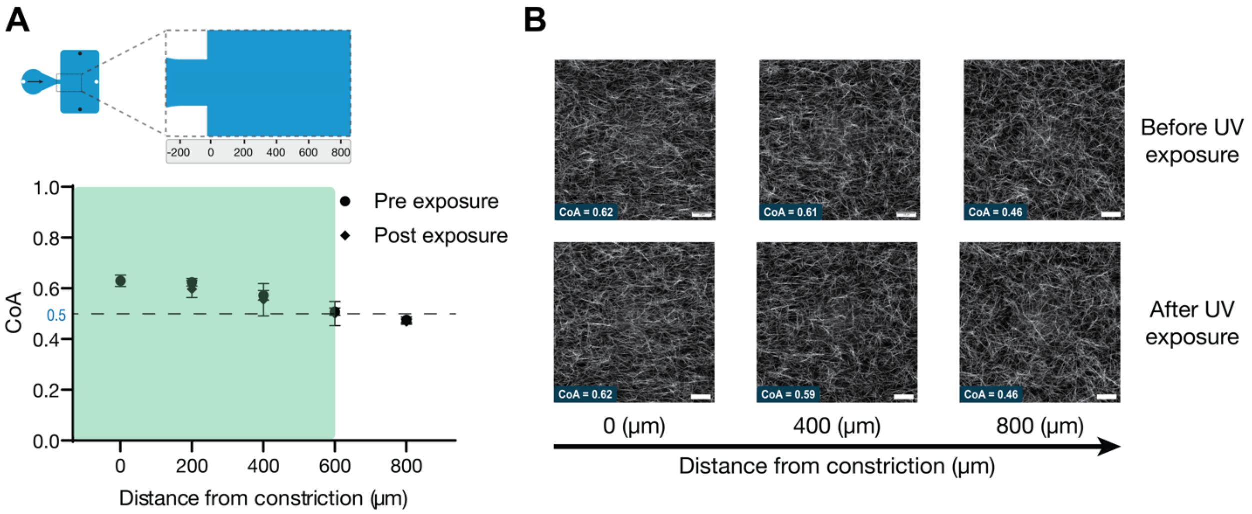

**Figure S7: A)** Shows the representative confocal images of COL1 fiber alignment before and after lifting the PDMS lid. **B)** Shows the graph of the change in CoA before and after lifting the PDMS channel, illustrating a minimal change in CoA (n=16).

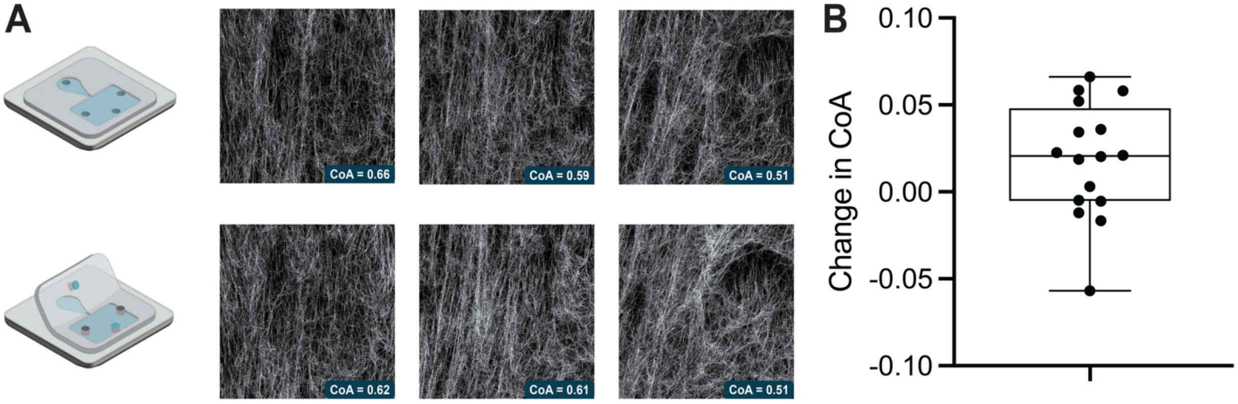


#### Deborah Number Calculation

De = $\lambda/\tau$

Where,

$\lambda$ = relaxation time = 0.0071 s [1]

$\tau$ = flow time scale = Length of constriction (mm) / Max velocity in constriction (mm/s)

References:


[1] A. Ahmed, I.M. Joshi, S. Larson, M. Mansouri, S. Gholizadeh, Z. Allahyari, F. Forouzandeh, D.A. Borkholder, T.R. Gaborski, V.V. Abhyankar, Microengineered 3D Collagen Gels with Independently Tunable Fiber Anisotropy and Directionality, Advanced Materials Technologies. n/a (n.d.) 2001186. https://doi.org/10.1002/admt.202001186.
